# Proteomic analysis of rice leaves following treatment of *Xanthomonas oryzae* pv. *oryzae* secreted cell wall degrading enzyme

**DOI:** 10.1101/2021.11.17.469048

**Authors:** Anirudh Kumar, Kamal Kumar Malukani, Ramya Pamidimukkala, Hitendra Kumar Patel, Ramesh V Sonti

## Abstract

Bacterial Blight (BB) disease caused by *Xanthomonas oryzae* pv. *oryzae* (*Xoo*), is one of the most devastating diseases in various rice cultivating countries. *Xoo* secretes a mixture of plant cell wall degrading enzymes (CWDEs) such as cellulases, lipases, xylanases, and proteases to degrade different components of the plant cell wall. LipA; a lipase/esterase, is one such *Xoo* secreted CWDE and is an important virulence factor of *Xoo*. Treatment of rice tissue with purified LipA induces immune responses. In this study, a LC-MS based proteomics study was performed to identify the differentially expressed proteins (DEPs) in rice following LipA treatment. A total of 212 proteins were identified in control and 201 proteins in LipA treated samples. There were 151 proteins common between control and treatment. Fold change analysis of these common proteins through SIEVE identified 26 upregulated and 49 downregulated proteins by at least ≥1.5 fold in the LipA treated sample. Pathway analysis indicated that many proteins related to redox regulation, photosynthesis, and translation are differentially expressed after LipA treatment. We also observed that some of the differentially expressed proteins contain translation regulatory elements that may regulate translation after LipA treatment. The comparison of proteomics data with previously performed transcriptome analysis indicated that different sets of genes and pathways are altered in both the analyses.

## Introduction

*Xanthomonas oryzae* pv. *oryzae* (*Xoo*), the causal agent of bacterial blight of rice secretes a battery of cell wall degrading enzymes such as cellulases, xylanases, and a lipase/esterase (LipA), to degrade different components of the rice cell wall (Rajeshwari et al., 2005, Jha et al., 2007). Cellulases and xylanases degrade polysaccharides component of the plant cell wall whereas LipA likely breaks the ester cross-links between the polysaccharide fibrils (Esquerré-Tugayé et al., 2000). These cell wall degrading enzymes have been proven to be important for virulence as mutation in any of these genes leads to virulence deficiency in *Xoo* (Rajeshwari et al., 2005, Jha et al., 2007). However, plant cell wall degrading enzymes are considered as double-edged swords as plants can recognize damage on the wall as a mark of infection and induce immune responses (Darvill and Albersheim, 1984, C A Ryan and Farmer, 1991). These immune responses include the production of secondary metabolites, reactive oxygen species (ROS) production, strengthening of the cell wall by callose deposition, and programmed cell death (PCD) (Dodds and Rathjen, 2010, Ghosh et al., 2019). Successful pathogens can cause disease only because they can suppress these host innate immune responses (Jha et al., 2005, Bent and Mackey, 2007).

Prior treatment of rice to purified *Xoo* secreted CWDEs induces resistance against subsequent infection of *Xoo* (Jha et al., 2007). Very little information is known, at the mechanistic level, about how plants mount defense responses following damage to their cell walls. To address this, a few microarray studies were performed following the treatment of rice tissue with *Xoo* secreted CWDEs (Malukani et al., 2020, Jha et al., 2010, Ranjan et al., 2015). However, sometimes these transcriptional changes do not directly reflect in changes at the proteome level (Liu et al., 2016). Poor correlation between mRNA abundance and translation was observed after treatment of Arabidopsis with elf18, an elicitor of immune response (Xu et al., 2017). The level of functional proteins is also regulated by translation, post-translation modification, and degradation. Therefore, an investigation of protein level is a prerequisite, and thus, this important aspect must be investigated via proteomics analyses.

In the last decade proteome analysis of plant samples has been mostly accomplished by the combination of 2-dimensional gel electrophoresis (2-DE) and mass spectrometry (MS). This technique has been used to study the effect of various biotic stresses in rice such as rice blast (Konishi et al., 2001), rice yellow mottle virus (Ventelon-Debout et al., 2004), and bacterial blight (Mahmood et al., 2006, Mahmood et al., 2009, Kumar et al., 2015). However, 2-DE suffers from certain shortcomings like ambiguity in the identification of multiple proteins present in a single spot, difficulties in identification of proteins at both extremes of pI range, natural variants, small proteins, and variation in extraction efficiency (Vissers et al., 2007). Although LC-MS-based isotope labeling provides greater accuracy, it also holds some disadvantages like high cost, procedural complexity, and potential artifacts. Therefore, label-free LC-MS quantification provides a better option to determine the relative abundance of protein in different samples based on measurement and comparison of MS signal intensities of peptide precursor ions (Wang et al., 2003, Silva et al., 2006, America et al., 2006).

In the current study, we have used a proteomics approach to identify rice proteins that are altered following treatment with LipA, a *Xoo* secreted cell wall degrading enzyme. Previously, high-resolution crystal structure of LipA has led to the identification of distinct ligand-binding modules (Aparna et al., 2009). Transcriptome analysis 12 h after LipA treatment indicated activation of the Jasmonic Acid (JA) pathway and further identification of various defense-related genes (Ranjan et al., 2015, Pillai et al., 2018, Kachewar et al., 2019). Transcriptome analysis after 2 h of LipA treatment led to the identification of a rice receptor kinase that likely perceived the damage caused by LipA (Malukani et al., 2020). The results of the current study indicate that a number of proteins that are associated with metabolism, translation, photosynthesis, redox status, and protein degradation are differentially expressed after LipA treatment. We further compared proteomics data with transcriptomic data and observed no clear correlation between them on gene-level or pathways level. We have also observed that the translation of some of these differentially expressed proteins might be regulated by upstream open reading frame (uORF) elements.

## Materials and Methods

### Plant growth conditions and LipA purification

Sterlized rice seeds were germinated in growth chamber and one week old seedlings were transferred to pots in green house with temperature maximum around 28ºC and minimum around 22ºC. LipA was purified from a culture supernatant of BXO2008, which is a LipA overexpression clone of *Xoo*. The LipA containing fractions were separated using cation exchange and size exclusion chromatography as described previously (Aparna et al., 2009). Quantification was made through NanoDrop spectrophotometer and its integrity and quality were examined by running on 12% SDS PAGE. The biological activity of purified LipA was tested using callose deposition and HR in rice leaves and programmed cell death in rice roots as described previously (Jha et al. 2007).

### LipA treatment, protein purification, and quantification

Rice leaves of 12-14 days old seedlings were infiltrated either with 100 μl of phosphate buffer (10 mM potassium phosphate, pH 6.6) (control) or 100 μl of purified LipA (0.5 mg/ml) (treatment) using 1.0 ml needleless syringes. After 2 h, around 100 leaf pieces covering the infiltrated zone from each treatment were harvested, frozen in liquid nitrogen, and stored in -80ºC.

Total leaf proteins were extracted as described by Parker et al. (2006) and Kumar et al. (2015) with little modifications (Parker et al., 2006, Kumar et al., 2015). Briefly, one gram of leaf tissue was homogenized to a fine powder and suspended in 5 ml of protein extraction buffer (0.5 M Tris-HCl pH 7.5, 0.7 M sucrose, 0.1 M KCl, 50 mM EDTA, 2% β-mercaptoethanol, and 1 mM PMSF). An equal volume of phenol saturated with Tris-HCl (pH 7.5) was added and mixed at 4ºC for 30 min followed by centrifugation at 6,000g for 20 min. The upper phenolic phase was collected and an equal volume of extraction buffer was added and the above step was repeated. Four volumes of 0.1M ammonium acetate in methanol was added and protein precipitation was allowed for 4 h at -20ºC. Subsequently, centrifugation was carried out at 10,000g for 30 min at 4ºC, and the pellet was washed three times with ice-cold methanol and acetone at 7,000g. Finally, dried pellets were solubilized in 250 μl of the rehydration solution [(8 M (w/v) urea, 2 M (w/v) thiourea, 4% (w/v) CHAPS, 30 mM DTT, 0.8% (v/v) immobilized pH gradient (IPG) buffer pH range 4–7 (GE Healthcare, Uppsala, Sweden)]. Finally, the concentration of protein was determined by the Bradford method (Bradford, 1976).

### Sample preparation and in-gel trypsin digestion for Nano-LC-MS/MS analysis

Approximately 100 μg protein samples were separated on 12% SDS PAGE at 100 V, stained in colloidal coomassie blue and washed subsequently. Gel slices were cut into small pieces and were washed with 50% acetonitrile (ACN) in 25 mM ammonium bicarbonate (NH_4_HCO_3_). Later, the gel pieces were reduced and alkylated by 10 mM DTT in 100 mM NH_4_HCO_3_ and 50 mM iodoacetamide in 100 mM NH_4_HCO_3_ and washed with 100 mM NH_4_HCO_3_ followed by dehydration in 50% ACN.

In-gel tryptic digestion was performed for LC-ESI-MS/MS Orbitrap Velos according to Li et al. (2012a) using 100 ng/μl trypsin (Promega, Madison, WI) for 16 h at 37ºC (Li et al., 2012). The trypsin-digested peptides were re-extracted by vortexing at room temperature (RT) with 0.1% trifluoroacetic acid (TFA) and 50% ACN in two steps of 60 min each. The peptide samples were desalted and purified with C18 ZipTip (Millipore ZipTips, micro-C18). The 1.0 μl of each sample was loaded on MALDI plate with 1.0 μl of the matrix. After MALDI confirmation, 10 μl of (0.1% formic acid +5% ACN/MQ) were added to each tube and centrifuged at 12000 rpm for 15 minutes followed by loading for LC-ESI-MS/MS.

### LC-MS/MS of trypsin digested proteins

In-gel digested peptide samples were analyzed on LTQ-Orbitrap Velos mass spectrometer (Thermo Scientific, Germany) interfaced with Easy-nLC II nanoflow liquid chromatography systems (Thermo Scientific, Odense, Southern Denmark). The peptide digests were reconstituted in solvent A (0.1% formic acid and fractionated on nanoflow LC system (Easy nLCTM II, Proxeon Biosystems, Odense, Denmark) using Bio Basic C18 Pico Frit nano-capillary column (75μ mX10cm: New objective, MA, USA) with a 90 min linear gradient 5-95% B [5% ACN with 0.1% formic acid (solvent A) and 95% ACN with 0.1% formic acid (solvent B)] at a flow rate of 300 nl/min and analyzed on LTQ Orbitrap Velos (Thermo Scientific, Waltham, MA, USA). 1.7kV was applied for ionization. Full scan MS with a mass window 350-1800 Da were acquired at1.7kV. Fragment ions were scanned in a low-pressure ion trap at a scan rate of 33,333 amu/s, a minimum threshold of ion selection for MS/MS was set at 500 counts. Ion accumulation times were set at 500 ms for MS and 25 ms for MS/MS. Activation time of 10 ms and q value of 0.25 was used.

### Identification of proteins

The MS/MS spectra of the multiple charged peptides obtained from all LC-MS were searched against the International Rice Genome Sequencing Project (IRGSP) database (Build 5) (http://rgp.dna.affrc.go.jp/E/IRGSP/Build5/build5.html) using SEQUEST search algorithms through Proteome Discoverer 1.5 Platform (Thermo Scientific, Bremen, Germany). The parameters used for data analysis included trypsin as the protease with a maximum of three missed cleavages. Carbamidomethylation of cysteine was specified as a fixed modification and oxidation of methionine as variable modifications. The minimum peptide length was specified to be 6 amino acids. Peptides were considered identified at a q value of 0.01 with a median mass measurement error of approximately 260 parts per billion. The MS/MS spectra were also examined manually for the proteins identified with single peptide hits. Proteins that had not shown three consistent b or y ions with good intensity were discarded from the protein list.

### Label-Free protein quantification

Peptide mixtures were subjected to nano-liquid chromatography coupled with MS and Bio Basic C18 Pico Frit nano capillary column (75μmX10cm: New objective, MA, USA). The analysis was carried out in a continuous acetonitrile gradient consisting of 5–45% B for 90 min (B = 95% acetonitrile, 0.5% acetic acid). The flow rate of 300 nl/min was maintained to elute peptides and for real-time ionization and peptide fragmentation at 1.7 kV. An enhanced FT-resolution spectrum (resolution = 60000) followed by the MS/MS spectra from the ten most intense parent ions were analyzed along the chromatographic run (90 min). All identifications were performed by Proteome Discoverer 1.4 software (Thermo-Fisher). Raw data files were processed and compared with SIEVE2 version 2.1 X86 (Thermo-Fisher).

For quantitative proteome analysis, six MS raw files from each pooled group were analyzed using SIEVE 2.1 software (Thermo Scientific, San Jose, CA, USA). Signal processing was performed in a total of 12 MS raw files. The SIEVE experimental workflow was defined as “Control Compare Trend Analysis” where one class of samples are compared to one or more other class of samples. The control samples were compared to each of the samples that were treated with LipA at 2 h time point. Coefficient correlation score values were acquired for each chromatogram and mean score values were calculated for each group. The values were in the range of 0.83 to 1.0 in control and, 0.78 to 0.91 in LipA treated sample.

The parameters used consisted of a frame m/z width of 1500 ppm and a retention time width of 1.75 min. A total of 21,635 MS2 scans were present in the files. Further, peak integration was performed for each frame and these values were used for statistical analysis. Subsequently, peptides were grouped into proteins and a protein ratio and p-value were calculated. Relative abundance of an individual protein from the control group was considered significantly different protein level when the values observed were <0.5 for decrease abundance or > 1.5 for increased abundance, and a p-value <0.05 as described.

### Pathway enrichment and ORF analysis

Pathway analysis was performed using the MAPMAN tool (http://mapman.gabipd.org/home) and g:Profiler online tool (https://biit.cs.ut.ee/gprofiler/gost) (Thimm et al., 2004, Raudvere et al., 2019). Proteins that were showing either upregulation or downregulation in at least two of the three sets were considered upregulated or downregulated respectively. Proteins that were unique to control (61) or downregulated (49) after LipA treatment were considered downregulated (Total-110). Similarly, proteins that were upregulated (26) after treatment or unique to treatment (50) were considered upregulated (Total 76). Analysis of the presence of an open reading frame was performed by the online tool uORFlight (http://uorflight.whu.edu.cn/home.html) (Niu et al., 2020). This was performed after converting RAP ID (obtained by proteomics) to rice MSU ID. Some of the RAP IDs did not have respective MSU ID, so this was performed with 71 upregulated (out of 76) and 107 downregulated (out of 110) proteins. Each gene that was differentially expressed on proteomics analysis was individually searched in uORFlight webpage tool and the list was assembled manually. All the Venn diagrams were created using the online tool Venny 2.0 (Oliveros, 2016).

## Results

### Nano-LC-MS/MS-based protein identification

Treatment of rice leaves with a lower concentration of LipA (0.1 mg/ml) induces callose deposition whereas treatment with a higher concentration of LipA (0.5 mg/ml) induces hypersensitive response (HR) like phenotype in rice leaves and programmed cell death in rice roots (**Supplemental Fig**.**1**). Since treatment of a higher concentration (0.5 mg/ml) of LipA leads to more robust defense response, we decided to use this concentration for our proteomics study. Nano-LC-MS/MS based analysis led to the identification of 212 common proteins in at least two sets of experiments out of three in control samples (buffer infiltration) whereas 201 proteins were common in at least two sets of experiments out of three in treatment samples (LipA infiltration) (**Fig. 1 A**). Further analysis revealed that there were 151 proteins common between controls and treatments, 61 were unique to controls and 50 were unique to treatments (**Fig. 1 A**). However, when fold change analysis was carried out using SIEVE with 151 common proteins, 26 proteins were found to be upregulated by a minimum of ≥ 1.5 fold change and 49 proteins were downregulated by at least ≤ -1.5 fold after 2 h of LipA treatment. The total upregulated and downregulated proteins are listed in **Table 1 and Table 2**. The PI of the entire set of differentially expressed proteins ranged from 4 to 11, the molecular weight ranged from 11 kDa to 111 kDa, and query coverage from 4% to 65%.

**Table 1:**
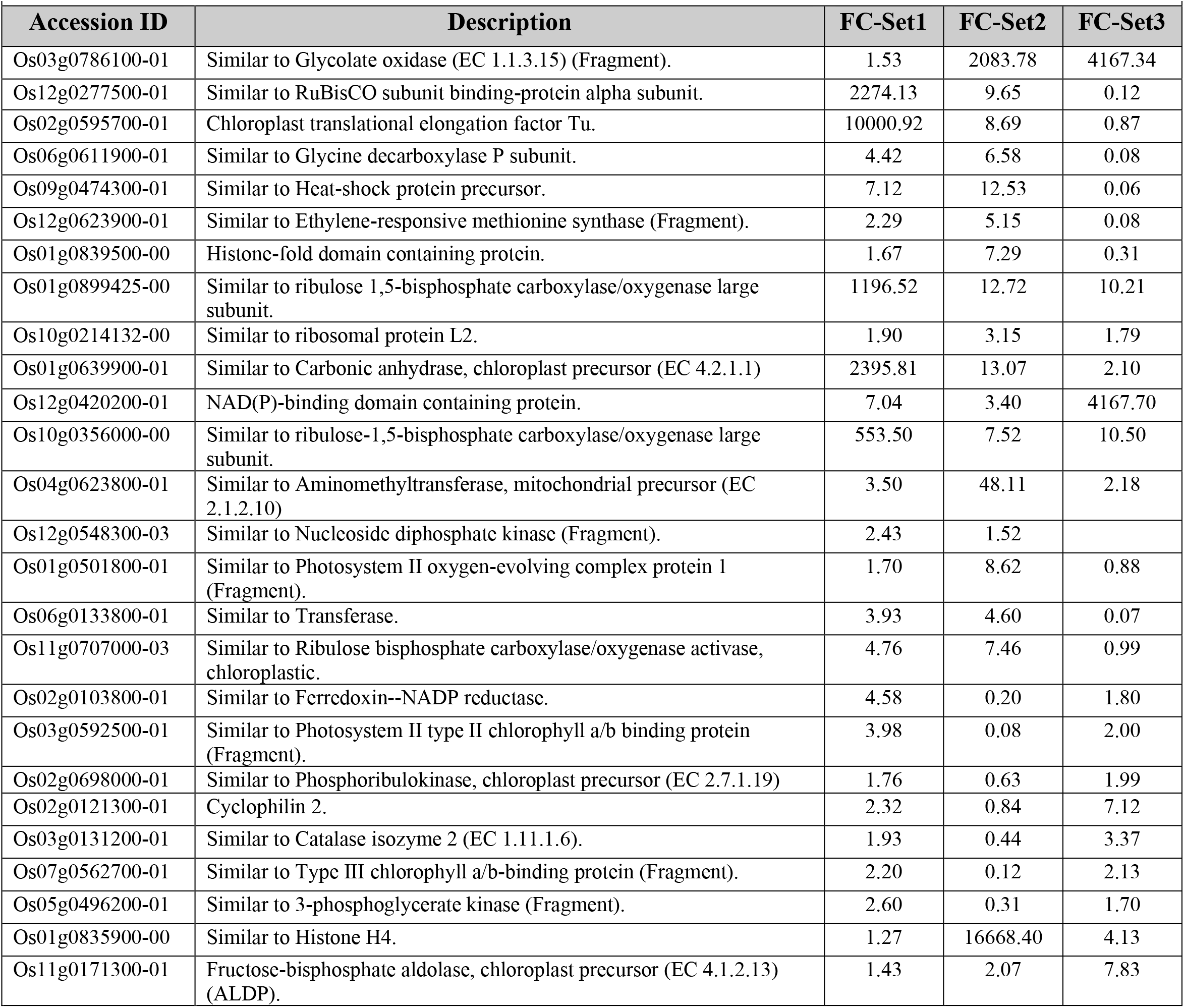
List of up-regulated proteins ≥ 1.5 fold change in at least 2 set upon LipA treatment, a cell wall degrading enzyme secreted by *Xanthomonas oryzae* pv. *oryzae*

| Accession ID | Description | FC-Set1 | FC-Set2 | FC-Set3 |
| --- | --- | --- | --- | --- |
| Os03g0786100-01 | Similar to Glycolate oxidase (EC 1.1.3.15) (Fragment). | 1.53 | 2083.78 | 4167.34 |
| Os12g0277500-01 | Similar to RuBisCO subunit binding-protein alpha subunit. | 2274.13 | 9.65 | 0.12 |
| Os02g0595700-01 | Chloroplast translational elongation factor Tu. | 10000.92 | 8.69 | 0.87 |
| Os06g0611900-01 | Similar to Glycine decarboxylase P subunit. | 4.42 | 6.58 | 0.08 |
| Os09g0474300-01 | Similar to Heat-shock protein precursor. | 7.12 | 12.53 | 0.06 |
| Os12g0623900-01 | Similar to Ethylene-responsive methionine synthase (Fragment). | 2.29 | 5.15 | 0.08 |
| Os01g0839500-00 | Histone-fold domain containing protein. | 1.67 | 7.29 | 0.31 |
| Os01g0899425-00 | Similar to ribulose 1,5-bisphosphate carboxylase/oxygenase large subunit. | 1196.52 | 12.72 | 10.21 |
| Os10g0214132-00 | Similar to ribosomal protein L2. | 1.90 | 3.15 | 1.79 |
| Os01g0639900-01 | Similar to Carbonic anhydrase, chloroplast precursor (EC 4.2.1.1) | 2395.81 | 13.07 | 2.10 |
| Os12g0420200-01 | NAD(P)-binding domain containing protein. | 7.04 | 3.40 | 4167.70 |
| Os10g0356000-00 | Similar to ribulose-1,5-bisphosphate carboxylase/oxygenase large subunit. | 553.50 | 7.52 | 10.50 |
| Os04g0623800-01 | Similar to Aminomethyltransferase, mitochondrial precursor (EC 2.1.2.10) | 3.50 | 48.11 | 2.18 |
| Os12g0548300-03 | Similar to Nucleoside diphosphate kinase (Fragment). | 2.43 | 1.52 |  |
| Os01g0501800-01 | Similar to Photosystem II oxygen-evolving complex protein 1 (Fragment). | 1.70 | 8.62 | 0.88 |
| Os06g0133800-01 | Similar to Transferase. | 3.93 | 4.60 | 0.07 |
| Os11g0707000-03 | Similar to Ribulose bisphosphate carboxylase/oxygenase activase, chloroplastic. | 4.76 | 7.46 | 0.99 |
| Os02g0103800-01 | Similar to Ferredoxin--NADP reductase. | 4.58 | 0.20 | 1.80 |
| Os03g0592500-01 | Similar to Photosystem II type II chlorophyll a/b binding protein (Fragment). | 3.98 | 0.08 | 2.00 |
| Os02g0698000-01 | Similar to Phosphoribulokinase, chloroplast precursor (EC 2.7.1.19) | 1.76 | 0.63 | 1.99 |
| Os02g0121300-01 | Cyclophilin 2. | 2.32 | 0.84 | 7.12 |
| Os03g0131200-01 | Similar to Catalase isozyme 2 (EC 1.11.1.6). | 1.93 | 0.44 | 3.37 |
| Os07g0562700-01 | Similar to Type III chlorophyll a/b-binding protein (Fragment). | 2.20 | 0.12 | 2.13 |
| Os05g0496200-01 | Similar to 3-phosphoglycerate kinase (Fragment). | 2.60 | 0.31 | 1.70 |
| Os01g0835900-00 | Similar to Histone H4. | 1.27 | 16668.40 | 4.13 |
| Os11g0171300-01 | Fructose-bisphosphate aldolase, chloroplast precursor (EC 4.1.2.13) (ALDP). | 1.43 | 2.07 | 7.83 |

**Table 2:** List of down regulated proteins ≤ 1.5 fold change in at least 2 set upon LipA treatment, a cell wall degrading enzyme secreted by *Xanthomonas oryzae* pv. *oryzae*.

| Accession ID | Description | FC-Set1 | FC-Set2 | FC-Set3 |
| --- | --- | --- | --- | --- |
| Os07g0176900-02 | Similar to ribose-5-phosphate isomerase. | 0.74 | 0.34 | 0.28 |
| Os03g0125000-01 | Similar to 50S ribosomal protein L5, chloroplast. | 0.68 | 0.68 | 0.38 |
| Os07g0558500-01 | Inositol phosphatase-like protein. | 0.59 | 0.56 | 0.62 |
| Os02g0192700-02 | Similar to Thioredoxin peroxidase. | 0.57 | 0.54 | 0.40 |
| Os01g0147900-01 | Triosephosphate isomerase, cytosolic (EC 5.3.1.1) (TIM) (Triose-phosphate isomerase). | 0.63 | 0.38 | 0.65 |
| Os08g0126300-01 | Similar to Glyceraldehyde-3-phosphate dehydrogenase (Fragment). | 0.58 | 0.28 | 1.15 |
| Os08g0460000-01 | Similar to Germin-like protein 1 precursor. | 0.42 | 0.63 | 0.46 |
| Os07g0556200-01 | Rieske [2Fe-2S] region domain containing protein. | 0.42 | 0.51 | 0.82 |
| Os02g0822600-01 | Similar to 50S ribosomal protein L9. | 0.73 | 0.69 | 0.63 |
| Os04g0486600-02 | Similar to Glyceralde-3-phosphate dehydrogenase. | 0.67 | 0.27 | 1.06 |
| Os08g0242700-01 | Similar to uridylyltransferase-related. | 0.31 | 0.22 | 0.27 |
| Os08g0266300-00 | Similar to predicted protein. | 0.71 | 0.30 | 0.20 |
| Os07g0694700-01 | L-ascorbate peroxidase. | 0.28 | 0.12 | 0.42 |
| Os12g0541500-02 | Similar to Chloroplast polypeptide of elongation factor Ts. | 0.71 | 0.73 | 0.46 |
| Os08g0561700-01 | Similar to Superoxide dismutase. | 0.47 | 0.66 | 0.18 |
| Os03g0379500-01 | Similar to 40S ribosomal protein S9. | 0.04 | 0.05 | 0.16 |
| Os06g0157000-01 | Esterase, SGNH hydrolase-type domain containing protein. | 0.40 | 0.16 |  |
| Os05g0388500-01 | Similar to 50S ribosomal protein L1. | 0.67 | 9.41 | 0.51 |
| Os02g0328300-01 | Phenol hydroxylase reductase family protein. | 0.34 | 12.67 | 0.17 |
| Os02g0115700-01 | Catalase isozyme A (EC 1.11.1.6) (CAT-A). | 0.40 | 2.52 | 0.27 |
| Os01g0265100-01 | Similar to ADP-ribosylation factor. | 0.60 | 4.83 | 0.58 |
| Os03g0284400-01 | Similar to Ribosomal protein L10-like. | 0.29 | 1.13 | 0.36 |
| Os08g0113100-02 | Similar to cDNA clone:J023041C07, full insert sequence. | 0.61 | 0.97 | 0.31 |
| Os08g0162600-01 | Rubredoxin-type Fe(Cys)<sub>4</sub> protein family protein. | 0.41 | 2.29 | 0.55 |
| Os07g0658400-02 | Similar to Fd-GOGAT protein (Fragment). | 1.03 | 0.41 | 0.33 |
| Os06g0725900-01 | Similar to Cell division protein ftsH homolog, chloroplast precursor (EC 3.4.24.-) (DS9). | 4.44 | 0.41 | 0.26 |
| Os03g0565200-01 | Haloacid dehalogenase-like hydrolase domain containing protein. | 1.25 | 0.66 | 0.75 |
| Os06g0114000-02 | Similar to 60 kDa chaperonin (Protein Cpn60) (groEL protein) (63 kDa stress protein). | 1.37 | 0.50 | 0.21 |
| Os08g0502700-03 | Similar to serine--glyoxylate aminotransferase. | 1.58 | 0.27 | 0.52 |
| Os02g0537700-01 | Similar to 2-cys peroxiredoxin BAS1, chloroplast precursor (EC 1.11.1.15) | 1.20 | 0.57 | 0.35 |
| Os07g0476900-01 | Thioredoxin domain 2 containing protein. | 1.23 | 0.15 | 0.58 |
| Os06g0669400-01 | Similar to Cell division protease ftsH homolog 2, chloroplastic. | 52.89 | 0.30 | 0.39 |
| Os05g0113900-01 | Similar to Histone H2A. | 2.72 | 0.54 | 0.37 |
| Os09g0485900-01 | Similar to 60S ribosomal protein L9 (Gibberellin-regulated protein GA). | 0.75 | 0.70 | 0.32 |
| Os03g0202200-01 | Porin-like protein. | 0.79 | 0.33 | 0.44 |
| Os05g0574400-01 | Similar to Malate dehydrogenase. | 0.80 | 0.28 | 0.67 |
| Os03g0356300-01 | Ribosomal protein L6 family protein. | 0.76 | 0.20 | 0.38 |
| Os03g0712700-01 | Similar to Phosphoglucomutase, cytoplasmic 2 (EC 5.4.2.2) | 1.34 | 0.50 | 0.29 |
| Os01g0205500-01 | Similar to 60S ribosomal protein L11-2 (L16). Splice isoform 2. | 0.76 | 0.45 | 0.44 |
| Os01g0869800-01 | Similar to Photosystem II subunit PsbS. | 1.61 | 0.22 | 0.54 |
| Os02g0750100-00 | Similar to ATP synthase. | 1.59 | 0.50 | 0.30 |
| Os08g0553800-01 | NAD(P)-binding domain containing protein. | 6.79 | 0.12 | 0.69 |
| Os06g0683200-01 | Similar to 50S ribosomal protein L24, chloroplast precursor (CL24). | 1.84 | 0.00 | 0.04 |
| Os01g0358400-01 | Similar to 40S ribosomal protein S4. | 0.92 | 0.74 | 0.20 |
| Os11g0703900-01 | Heat shock protein 70. | 1.16 | 0.71 | 0.08 |
| Os02g0115900-01 | Endosperm luminal binding protein. | 0.99 | 0.51 | 0.13 |
| Os02g0603800-01 | Similar to Isoprenoid biosynthesis-like protein (Fragment). | 1.64 | 0.51 | 0.17 |
| Os07g0558400-01 | Similar to Chlorophyll a/b-binding protein CP29 precursor. | 0.76 | 0.25 | 0.69 |
| Os12g0625000-01 | Similar to Cysteine synthase. | 1.93 | 0.56 | 0.70 |

**Figure 1:**
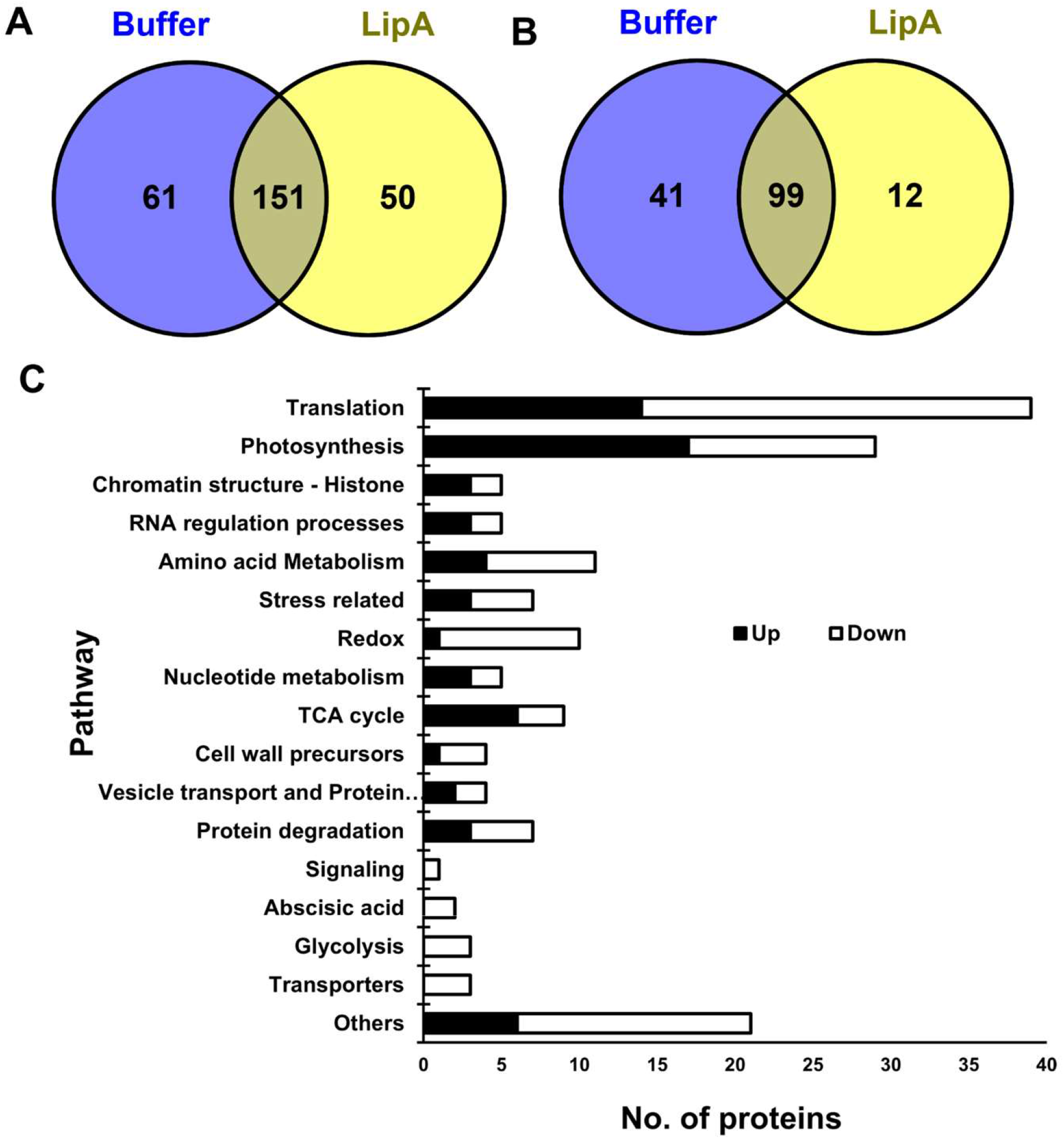
Proteins and pathways observed to be altered by proteomics analysis. (A) Venn diagram indicating the number of proteins observed in at least two of the three sets of treatment (0.5 mg/ml LipA) or control (10 mM potassium phosphate buffer). (B) Venn diagram indicating the number of proteins observed in all three sets of experiments. (C) Pathways altered in proteomics with genes observed in any two sets of experiments. Pathway analysis was done by the MAPMAN tool.

Subsequently, we selected those proteins which were present in all three sets of experiments. There were 140 proteins present in all three sets of control and 111 proteins were present in all three sets of treatments. Comparison of these subsets indicated that there were 99 proteins common in both control and treatments whereas 41 proteins were unique to control and 12 were unique to treatments (**Fig. 1 B**). When these 99 proteins (common in controls and treatments) were studied for fold change, 23 were found to be up-regulated (≥1.5 fold change) and 31 were down-regulated (≤ -1.5 fold change) in at least two out of three sets of experiments. The PI of the entire set of differentially expressed protein ranged from 4 to 11, molecular weight ranged from 11 kDa to 111 kDa, and query coverage from 4% to 65%.

### Pathway analysis of differentially expressed proteins

Pathway analysis using the MAPMAN tool indicated several proteins related to various pathways were differentially expressed after treatment. The proteins involved in redox regulation, translation, transporters, cell wall precursors, glycolysis, abscisic acid metabolism, and amino acid metabolism were more in the downregulated set of proteins. On the other hand, proteins involved in the Tricarboxylic acid cycle (TCA cycle) and photosynthesis were more in the upregulated set of proteins. Some proteins involved in photosynthesis, protein degradation, nucleotide metabolism, stress responses, RNA regulation and processing, and chromatin structure (histone) were also differentially expressed but no significant pattern was observed in those pathways (**Fig. 1 C**). Interestingly only 1 out of 7 differentially expressed proteins involved in stress responses are listed as being involved in biotic stress while the other 6 are listed as being involved in abiotic stress responses. Pathway analysis using g:Profiler tool provided more information about enriched pathways (**Supplemental Fig. 2**). Pathways involved in molecular functions indicated that proteins involved in antioxidant activity, pore complexes, and RNA binding are significantly downregulated whereas proteins involved in the oxygen-evolving complex are significantly upregulated (**Supplemental Fig. 2 A**). Similarly, KEGG analysis indicated proteins involved in the TCA cycle, folic acid, and amino acid synthesis were upregulated. On the other hand, proteins involved in the metabolism of glycine, serine, and threonine amino acids are significantly downregulated (**Supplemental Fig. 2 B**).

### Comparison between transcriptomics and proteomics data

We have previously performed transcriptome analysis after 2 h and 12 h of LipA treatment in rice (Malukani et al., 2020, Ranjan et al., 2015). This led to the identification of 78 and 2453 genes differentially expressed 2 h or 12 h respectively after following LipA treatment (FC ≥ 1.5). Comparison between transcriptomics and proteomics data indicates that none of the genes observed to be differentially altered after 2 h of LipA treatment were observed in our proteomics data performed at the same time point **(Fig. 2 A)**. We hypothesized that maybe some of these genes might be non-significantly upregulated in the microarray and filtered out by our criteria of ± 1.5 fold change or p<0.05. With this in mind, we examined the expression of these genes in unfiltered raw data and observed that none of them show difference in transcript levels **(Supplemental Table 1)**. We had also performed transcriptomics after 12 h of LipA treatment (Ranjan et al., 2015). We observed 52 genes are commonly present in microarray performed after 12 h of LipA treatment and MS/MS performed after 2 h of LipA treatment **(Fig. 2 A)**. Most of these 52 common genes (49) are downregulated in microarray analysis. While in MS/MS analysis, 18 genes are upregulated and 34 are downregulated. Overall, a comparison between microarray and MS/MS data indicates a poor correlation.

**Figure 2:**
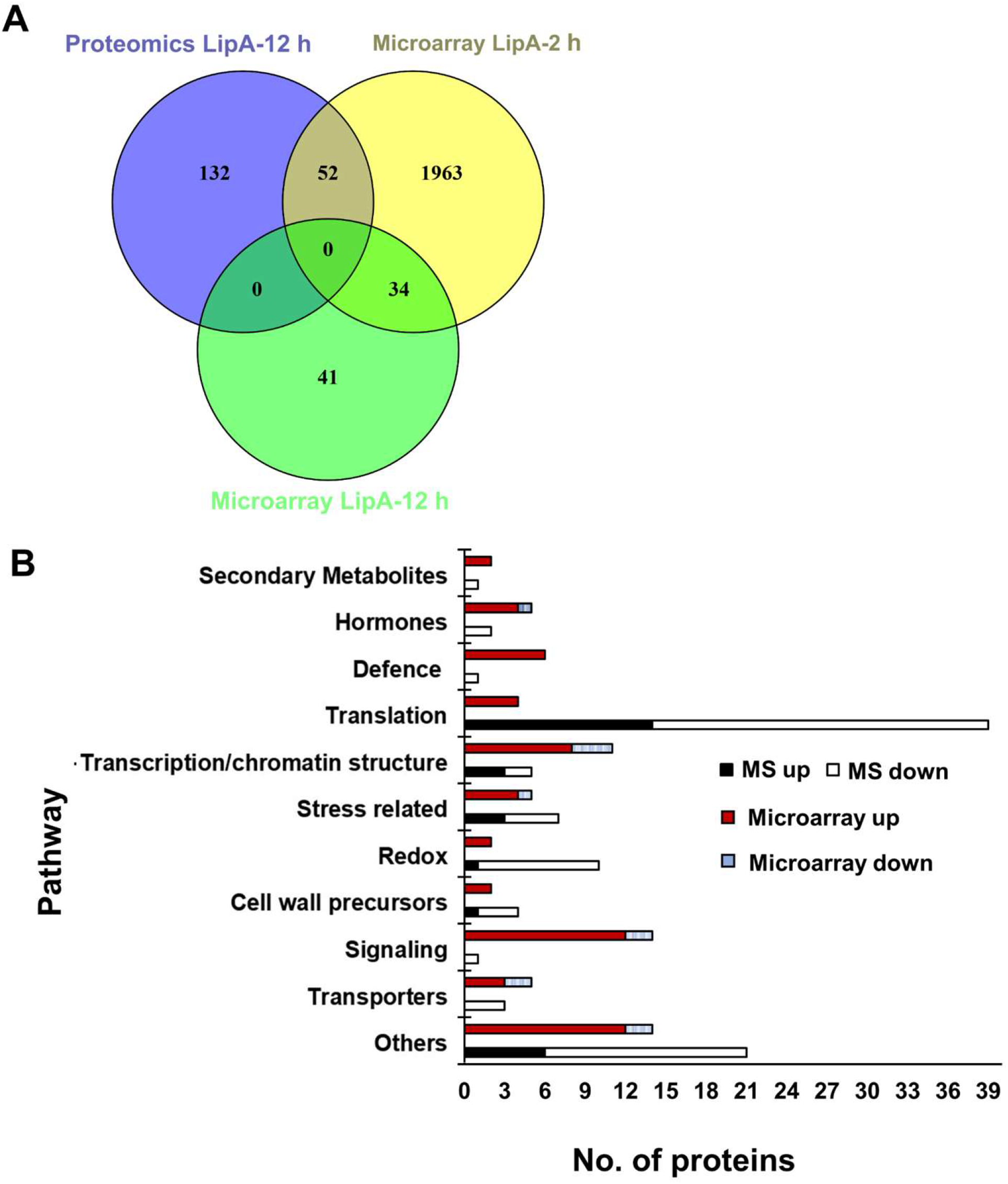
Comparison of microarray and proteomics data performed after treatment of 0.5 mg/ml LipA. (A) Venn diagram indicating the number of differentially expressed genes/proteins in microarray analysis and proteomic analysis. (B) Altered pathways common between transcriptomics and proteomics after 2 h of LipA treatment as observed by MAPMAN tool.

To test if similar pathways are enriched in transcriptomics and proteomics data, we compared MAPMAN analysis data of enriched proteins with the previously published data of microarray performed after 2 h of LipA treatment (Malukani et al., 2020) **(Fig. 2 B)**. The most striking difference was in genes involved in redox reactions. While microarray analysis indicated 2 genes being upregulated and 1 downregulated in redox reactions, proteomics indicated 9 proteins being downregulated while only 1 was upregulated. The genes involved in stress responses were upregulated in microarray, whereas they showed mixed profiles in the proteomics analysis. Similarly, while 4 genes related to protein synthesis (translation) and turnover were upregulated in microarray analysis, proteomics analysis indicated that 39 such proteins were differentially expressed with many of them being downregulated. While microarray analysis indicated upregulation of GA pathway-related genes, proteomics indicated downregulation of ABA pathway-related proteins. Many of the upregulated genes in microarray belong to the category of signal transduction with many of them being upregulated. Only one signal transduction protein (a 14-3-3) was observed in proteomics analysis and this was downregulated. The microarray data also indicated the upregulation of three receptor kinases which was not observed in proteomics. Three secondary metabolite related genes were upregulated in microarray analysis while only one secondary metabolite-associated protein was differentially expressed in proteomics and it was downregulated. A few categories observed only in proteomics analysis include the TCA cycle, glycolysis, amino acid metabolism, ABA synthesis, protein degradation, protein targeting, RNA regulation, chromatin structure, and photosynthesis.

### Analysis of open reading frame (ORF)

Variation at the protein level without change in the mRNA level could be because of multiple reasons. Two major reasons for this include translation regulation and protein degradation. Our pathway analysis already indicates differential expression of some translation-related proteins as well as proteins involved in protein degradation **(Fig. 2 A)**. To test if any of the differentially expressed proteins also contain translation regulator elements, we looked for the presence of an open reading frame (ORF) in the DNA sequence of these genes. We observed 13 of the unique 71 upregulate proteins and 25 of the 107 unique downregulated proteins contain regulatory ORF (upstream ORF and major ORF) (**Supplemental Table 2**). This indicates translation of at least some of the altered proteins might be regulated by respective ORF.

## Discussion

BB caused by *Xoo* is a serious threat to rice production globally. *Xoo* secretes a battery of enzymes to hydrolyze the plant cell wall. This includes a lipase/esterase (LipA). The LipA enzyme is important for virulence of *Xoo*, and might reduce the rigidity of the cell wall likely by disruption of the ester cross-links between the polysaccharide fibrils (Aparna et al., 2009). The results presented here represent the proteomic changes in plant cells that are undergoing an innate immune response induced upon cell wall damage caused by LipA in rice. We also compared this proteomics data with microarray data performed previously after the same treatment. We observed different sets of genes and pathways are differentially expressed in microarray and proteomics. Poor correlation like this between mRNA abundance and translation level is already reported in some studies including in rice (Liu et al., 2016, Peng et al., 2015).

The results indicate that there is a clear downregulation of ROS-related proteins. These mainly include ROS scavenging enzymes such as catalases, superoxide dismutase, ascorbate peroxidases, and thioredoxins (Huang et al., 2019, Dvořák et al., 2021). Such downregulation could promote the accumulation of ROS which can lead to immune responses (Dvořák et al., 2021). The induction of immune responses also reduces photosynthesis in plants which also helps in the accumulation of ROS (Su et al., 2018, Yang et al., 2021). This could be the reason many photosynthesis-related proteins are altered after LipA treatment. The third category where many proteins are differentially expressed is translation. This includes many ribosomal proteins subunits some of which are upregulated while some are downregulated. A few recent studies indicate the role of ribosomal proteins in plant immune responses (Ramu et al., 2020, Xu et al., 2017).

When we compared the proteomics data with microarray data, we observed there is hardly any overlap between those two data sets. The first observation was while most of the transcriptionally altered genes were upregulated, many of the differentially regulated proteins identified through proteomics were downregulated. There also seems to be a poor correlation in altered pathways observed by microarray and proteomics. We observed many pathways which were altered in proteomics analysis but not altered in transcriptomics. It is possible that this might be because of a smaller number of genes altered in transcriptomics than in proteomics. But at the same time, some categories such as transcription factors, signaling-related genes including receptor kinases, and hormone-related genes are more represented in transcriptome analysis. These results indicate that although the application of a single technology can provide insights into how immune responses are elaborated, the use of multiple omics-technologies can lead to a better understanding of the plant immune system which in turn can lead to more possibilities for translation in the field.

## Supporting information

Fold change and p-value of differentially expressed proteins in microarray data performed after 2hr of LipA treatment.

Analysis of uORF using uORFlight in DNA sequence of differentially expressed proteins after LipA treatment

## Abbreviations

BB: Bacterial Blight
PCD: Programmed Cell Death
*Xoo*: *Xanthomonas oryzae* pv. *oryzae*
PAMPs: Pathogen-Associated Molecular Patterns
MS: Mass Spectrometry
LC-MS: Liquid Chromatography-Mass Spectrometry
HR: Hypersensitive Response
DEPs: Differentially expressed proteins

## Figure legends

**Supplemental Figure 1:**
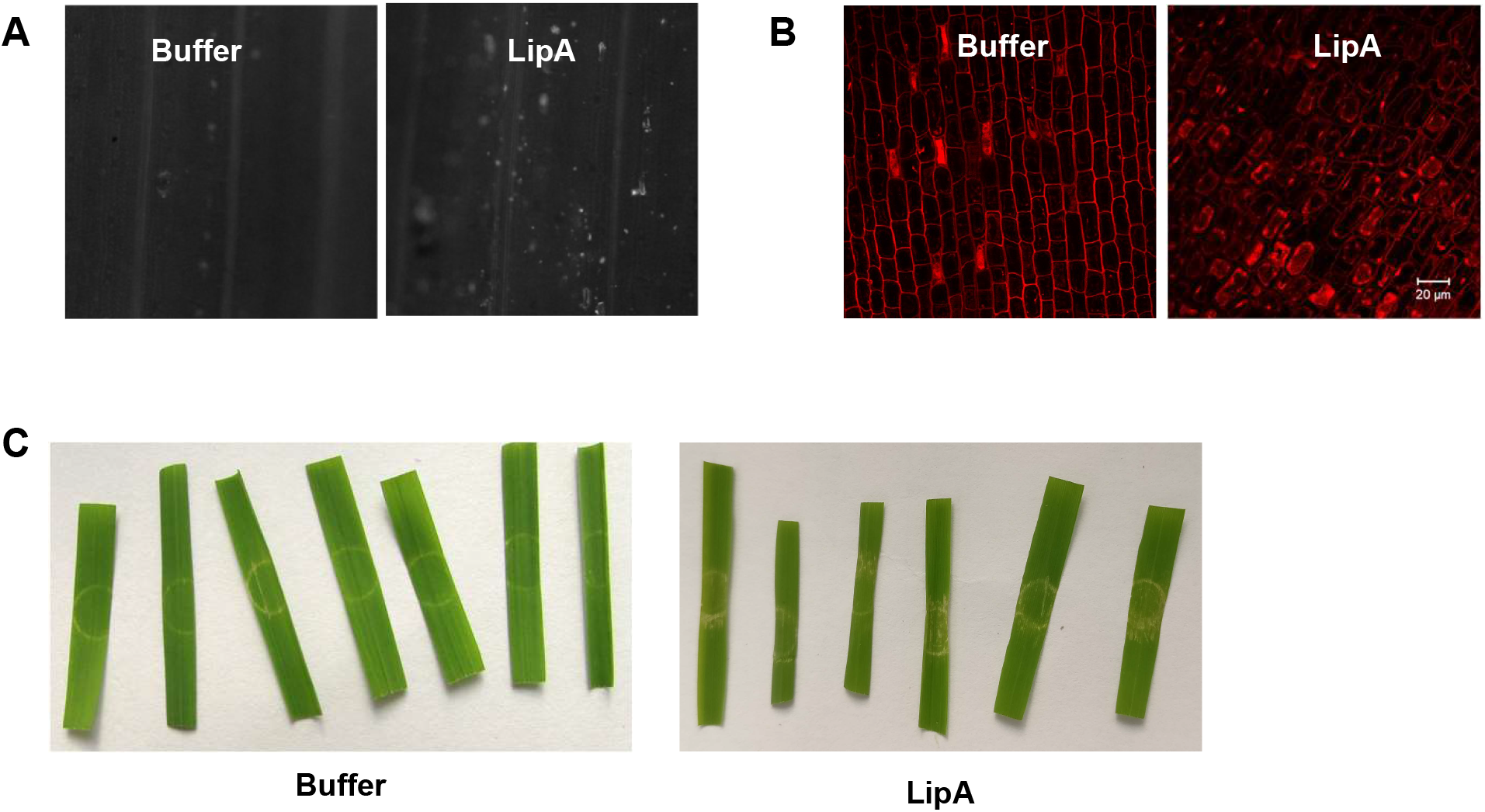
LipA treatment induces rice immune responses. (A) Treatment of 0.1 mg/ml of LipA activates callose deposition in rice leaves. (B) Treatment of rice roots with 0.5 mg/ml LipA induces programmed cell death. (C) Infiltration of rice leaves with 0.5 mg/ml LipA induces HR like phenotype.

**Supplemental Figure 2:**
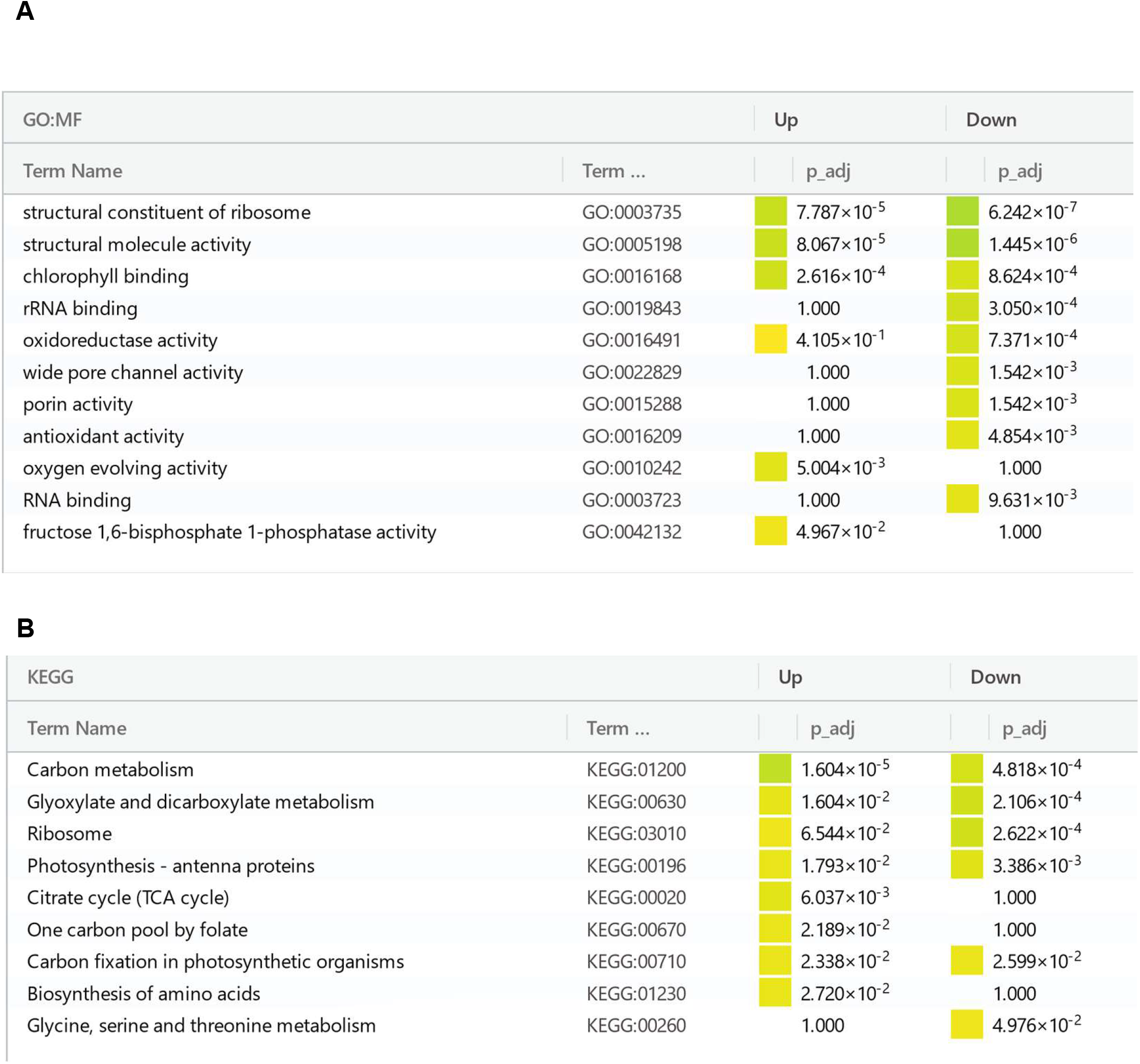
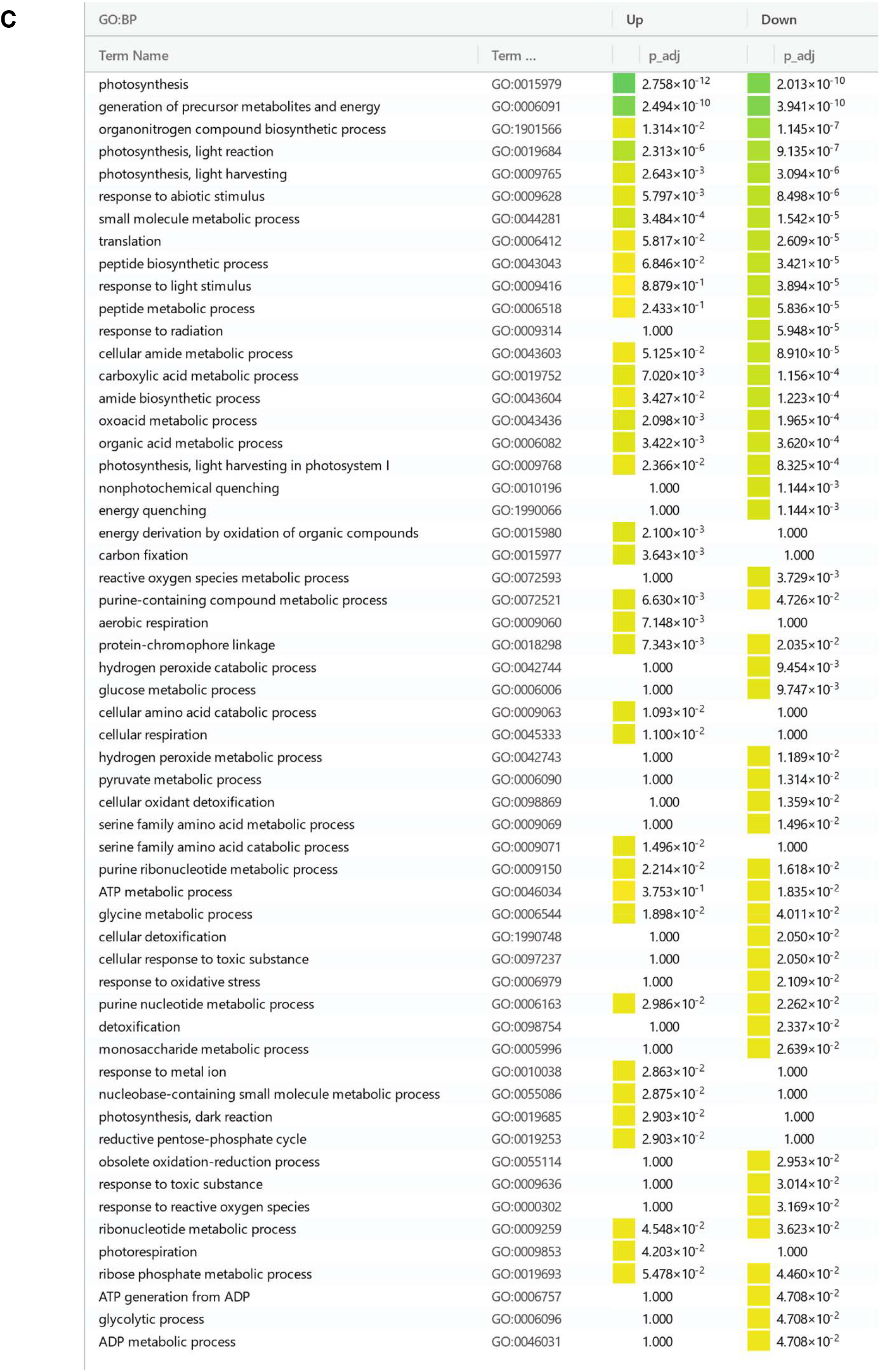
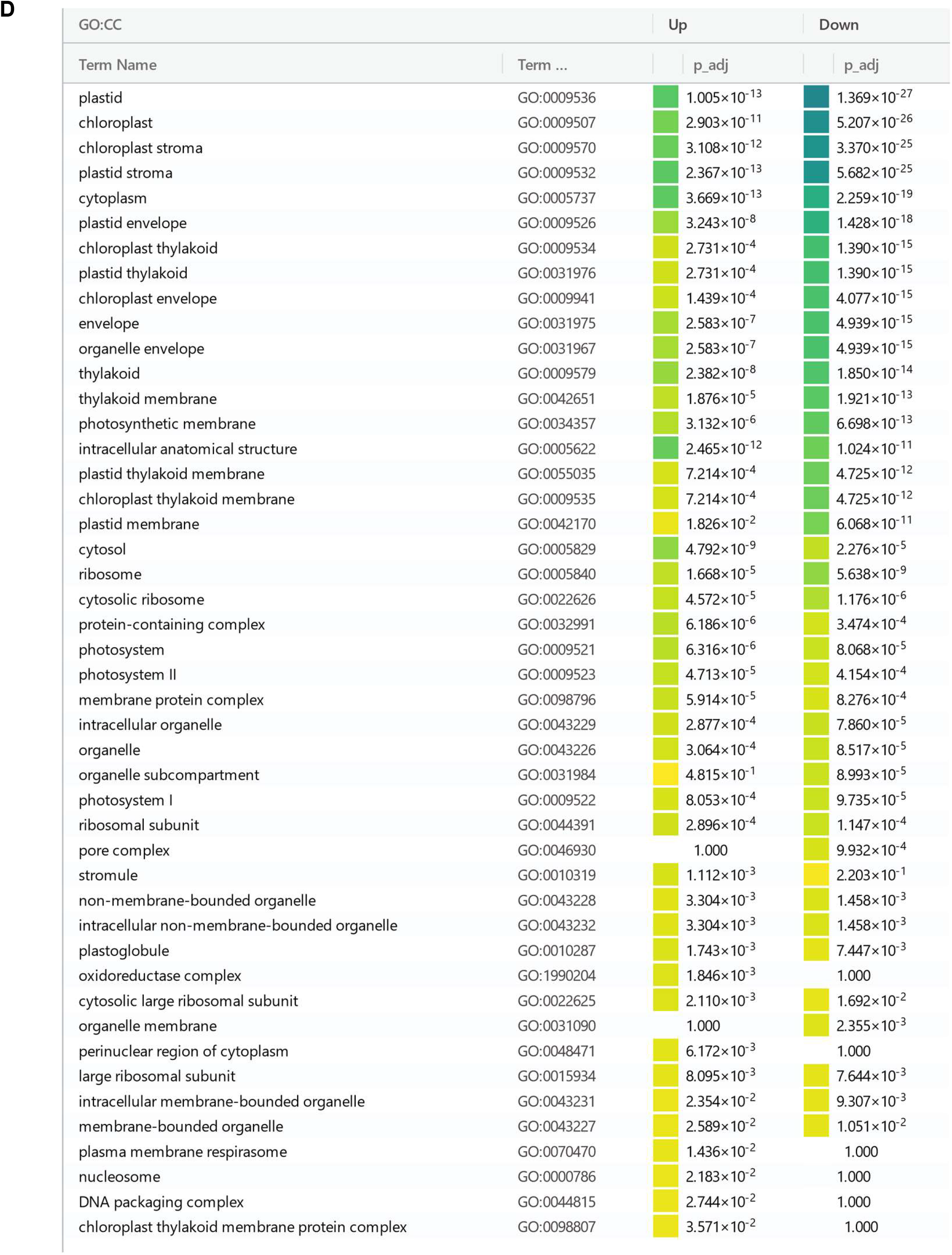
Pathways observed to be enriched in differentially expressed proteins. The analysis was performed using g:Profiler tool and the images were exported from the same tool The categories include molecular functions (MF) (A), Kyoto Encyclopedia of Genes and Genomes (KEGG) pathways (B), biological processes (BP) (C), and cellular components (CC) (D).

**Supplemental Table 1:** Fold change and p-value of differentially expressed proteins in microarray data performed previously after 2 h of LipA treatment.

**Supplemental Table 2:** Analysis of uORF using uORFlight in DNA sequence of differentially expressed proteins after LipA treatment.

